# Physiological Magnesium Concentrations Increase Fidelity of Diverse Reverse Transcriptases from HIV-1, HIV-2, and Foamy Virus, but not MuLV or AMV

**DOI:** 10.1101/2021.08.05.455312

**Authors:** Ruofan Wang, Ashton T. Belew, Vasudevan Achuthan, Najib El Sayed, Jeffrey J. DeStefano

## Abstract

Reverse transcriptases (RTs) are typically assayed in vitro using optimized Mg^2+^ concentrations (~ 5-10 mM) that are several-fold higher than physiological cellular free Mg^2+^ (~ 0.5 mM). Analysis of fidelity using *lacZα*-based α-complementation assays showed that tested HIV RTs, including HIV-1 from subtype B (HXB2-derived), HIV-2, subtype A/E, and several drug-resistant HXB2 derivatives all showed significantly higher fidelity using physiological Mg^2+^. This also occurred with prototype foamy virus (PFV) RT. In contrast, Moloney murine leukemia virus (MuLV) and avian myeloblastosis virus (AMV) RTs demonstrated equivalent fidelity in both low and high Mg^2+^. In 0.5 mM Mg^2+^, all RTs demonstrated ≈ equal fidelity, except for PFV RT which showed higher fidelity. A Next Generation Sequencing (NGS) approach that used barcoding to accurately determine mutation rates and profiles was used to examine the types of mutations made by HIV-1 (subtype B, wild type) in low (0.5 mM) and high (6 mM) Mg^2+^ with DNA or RNA that coded for *lacZα*. Unlike the α-complementation assay, which is dependent on LacZα activity, the NGS assay scores mutations at all positions and of every type. A ~ 4-fold increase in substitution mutations was observed in high Mg^2+^. The general trend was an exacerbation in high Mg^2+^ of more common mutation in low Mg^2+^, rather than the creation of new mutation hotspots. These findings help explain why HIV RT displays lower fidelity in vitro (with high Mg^2+^ concentrations) than other RTs (e.g., MuLV and AMV), yet cellular fidelity for these viruses is comparable.

## Introduction

Retroviral reverse transcriptases (RT) possess RNA- and DNA-dependent DNA polymerase activity and RNase H activity [1]. Magnesium (Mg^2+^), the most abundant divalent cation in the cell, functions as the physiological co-factor for both activities. Each of the two physically separated active sites is proposed to contain two divalent cation binding sites [2–8].

The enzymatic activities of purified HIV RT have typically been investigated in vitro using ~ 5-10 mM Mg^2+^, 25-100 μM dNTPs, and about 80 mM KCl or NaCl. These values approximate the optimal activity levels determined on homopolymeric templates for HIV RT polymerase and RNase H activities [9–12]. However, they are not representative of physiological levels of these components. This is often the case for general enzymatic analysis in vitro which is typically performed under optimized conditions with only required components, in part because it is not possible to completely mimic complex physiological conditions in vitro. Although the use of non-physiological levels of some components may not significancy effect enzyme properties, the concentration of Mg^2+^ can have profound effects on HIV RT DNA synthesis, drug interaction, and fidelity. While the total Mg^2+^ concentration in cells is high (typically 10 mM or more [13–15], most is sequestered by nucleotides and other complex anions, and “free” Mg^2+^ in lymphocytes and several other cell types has typically been found to be ~ 0.5 mM [13, 14, 16–21]. RNA-directed ssDNA synthesis reactions performed with HIV-1 RT with low Mg^2+^ lead to more efficient ssDNA synthesis, despite modestly decreased overall activity [22]. In in vitro reactions, many common nucleoside reverse transcriptase inhibitors (NRTIs) are less effective in lower Mg^2+^ while non-nucleoside reverse transcriptase inhibitors (NNRTIs) are more effective [23]. Finally, the fidelity of HIV RT, while independent of dNTP and KCl concentrations, is higher at lower, more physiological Mg^2+^ concentrations [24, 25]. These findings offer a possible explanation for why HIV RT displays lower fidelity in vitro (with high Mg^2+^ concentrations) than other reverse transcriptases (e.g., Moloney murine leukemia virus (MuLV) and avian myeloblastosis virus (AMV)), yet cellular fidelity for these viruses is comparable [26, 27].

In this report we examined the fidelity of HIV-1 RT wild type (wt) and several common drug-resistant mutants in high (6 mM) and low (0.5 mM) Mg^2+^ using a frequently employed *lacZα*-based α-complementation assay, to see if resistance mutations altered the effect of Mg^2+^ on RT fidelity. In addition, HIV-2, protype foamy virus (PFV), MuLV, and AMV RTs were also tested. A single strand consensus sequencing (SSCS) Next Generation Sequencing (NGS) analysis was also performed with a *lacZα* template using wt HIV-1 RT with high and low Mg^2+^. This produced a comprehensive mutation profile that, unlike α-complementation which can only detect mutations that decrease *lacZα* activity, includes all nucleotide sites on the *lacZα* gene. The results are compared to other RT fidelity assays conducted in vitro and in cells.

## Results

### System used to test the fidelity of RTs

The plasmid-based system used for in vitro fidelity analysis was similar to a previous system used in the lab [25]. Construction of the new plasmid (pBS∇EcoRV_567_) for this system is described in Materials and Methods and the region of the plasmid coding for the *lacZα* gene is shown in Fig. 1 along with the basic design of the system. This new system evaluates fidelity over a longer region of th*e lacZα* gene that matches up better with others used in the literature [28, 29]. The restriction sites used to insert RT-derived DNA include an EcoRI site in the plasmid’s MCS and an inserted EcoRV site. Unlike the original system, both sites are outside the *lacZα* gene region, minimizing the potential for white or faint blue colonies resulting from ligation artifacts, although deletion or insertion errors at the EcoRI site could still lead to frameshifts in *lacZα*. This system also truncates the *lacZα* gene relative to the previous by adding double stop codons before the EcoRV site. Background error rates for the system were assessed by cleaving pBS∇EcoRV_567_ with EcoRV and EcoRI, isolating the insert, and reinserting it into plasmids prepared for fidelity analysis, then transforming bacteria in α-complementation assays (see Methods). This approach produced ~ 1 white or faint blue colony per 1000 total colonies (plasmid preparations that produced significantly higher proportions of white and faint blue colonies were discarded). Note that this number did not change significantly compared to inserting DNA produced with Q5 DNA polymerase which was the control for the fidelity assays (Table S1, 16/18,371 and 10/11,078, for experiments started with RNA and DNA templates, respectively). This is likely due to the very high fidelity of Q5 DNA polymerase [30]. For a detailed explanation of the sources of mutations in the RNA and DNA templated versions of this assay see “Sources of mutations in the α*-* complementation assays” under Material and Methods.

**Figure 1.**
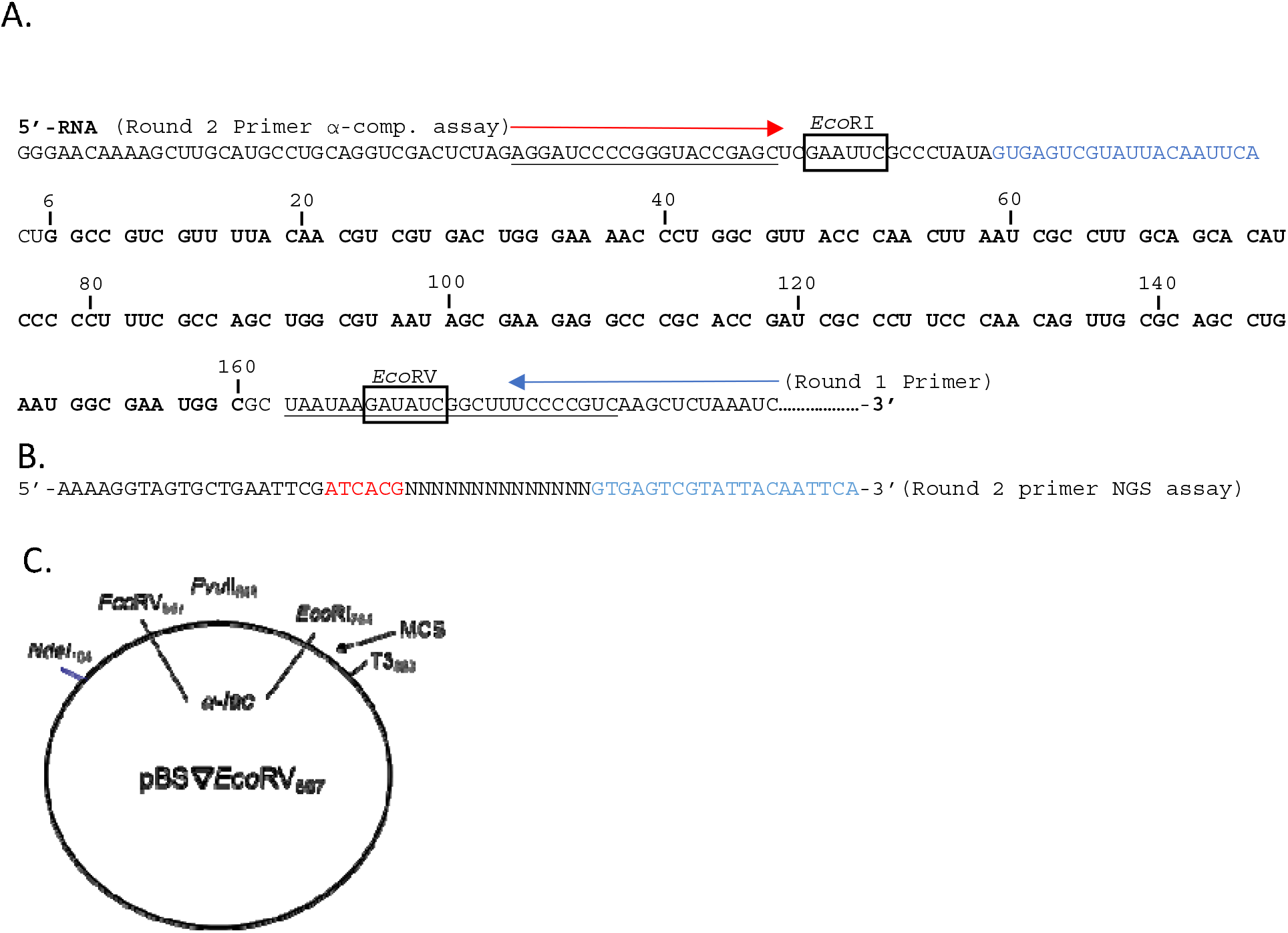
Constructs used to examine reverse transcriptase fidelity. (A) The relevant region of the RNA template made from T3 RNA polymerase is shown. A similar region was used for the DNA templated assay (see Methods). Numbering is only shown in the area of the *lacZα* gene and corresponds to the number used by Abram et al. (29). Codons for producing the LacZα peptide are in 3 base sets. The region of the sequence scored for mutations in the NGS assay are bolded. The α-complementation assay scored the region between the EcoRI and EcoRV sites. Sites of round 1 and 2 primers for the 2 rounds of RT synthesis in the RNA templated assays are depicted by arrows. The DNA template assay, which had only a single round of RT synthesis, used only the round 2 primer. (B) The sequence of the indexing primer for the NGS assay is shown. This primer was used as the round 2 primer in the RNA templated NGS assay and the round 1 primer (only round) in the DNA templated NGS assay. The primer contains a 14-nucleotide random region (N) that was used for indexing (see Methods) and a 6-nucleotide barcode (in red) used to identify the particular condition. The region in blue corresponds to the region shown in the same color in panel A. (C) Plasmid construct used in assays. Plasmid pBS∇EcoRV_567_ was constructed as described under Methods. Numbering is based on the parent plasmid, pBSM13+.

### With the exception of MuLV and AMV RTs, all RTs showed significantly higher mutation levels with 6 vs. 0.5 mM Mg^2+^ in *α*-complementation assays

Detailed results for analyses with several RTs are shown in Table S1, and the relative mutation frequencies (relative to the mutation frequency for wt HIV RT at 0.5 mM Mg^2+^) are shown in Fig. 2. For wt HIV RT with the RNA templated system, the analysis was performed at several Mg^2+^ concentrations, while other RTs were with 0.5 and 6, or 0.5, 6, and 12 mM Mg^2+^ only (all free Mg^2+^ concentrations). In the RNA templated system, mutation frequencies were typically 2-3-fold higher with 6 vs. 0.5 mM Mg^2+^ and increasing to 12 mM did not result in a significant increase above 6 mM. Using *p*<0.05 as a cutoff, all HIV enzymes, including HIV-2 RT, showed significantly greater mutation frequencies at 6 vs. 0.5 mM Mg^2+^ (Table S1, 2^nd^ column from right). In general, mutation frequencies of drug resistant mutants did not differ significantly from wt HIV RT under that same Mg^2+^ conditions (Table S1, last column on right). Two notable exceptions were K65R and M184V. At 6 mM Mg^2+^, the former was significantly lower while the latter was significantly higher than wt. The K65R results are consistent with previous results demonstrating that this mutation results in higher fidelity [31]. Like HIV RT, PFV RT was also less accurate in higher Mg^2+^, however, this enzyme showed significantly higher fidelity than HIV RT in both 0.5 and 6 mM Mg^2+^. We note that unlike our observations, studies from others did not find that PFV RT was more accurate than HIV RT [32]. Finally, MuLV and AMV RTs behaved differently than other RTs as neither showed a statistically significant difference in mutation frequencies in 0.5 vs. 6 mM Mg^2+^, and only MuLV RT at 6 mM Mg^2+^ was significantly different than HIV RT wt under that condition, demonstrating greater fidelity.

**Figure 2.**
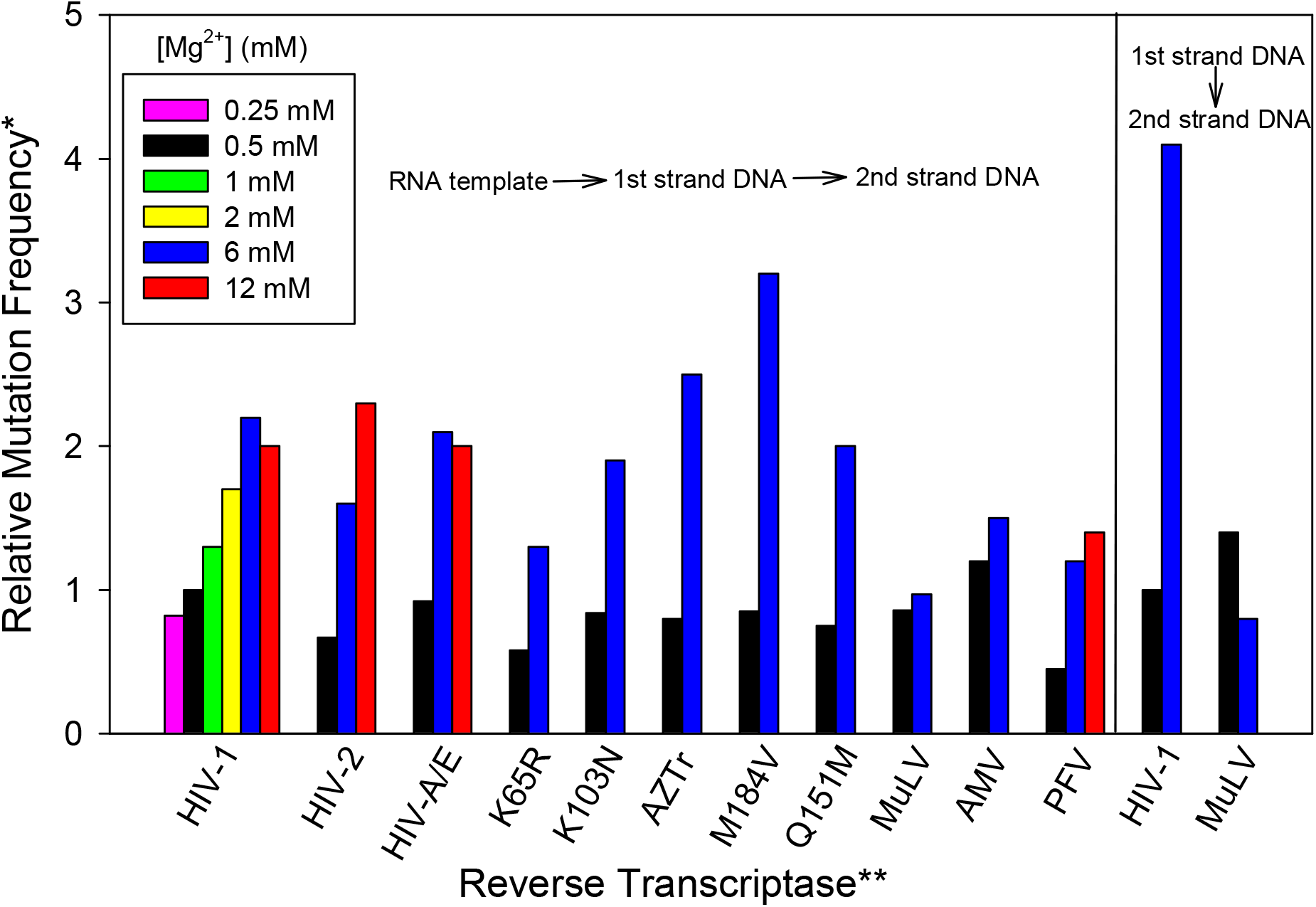
Relative fidelity of various reverse transcriptases (RT) in different Mg^2+^ concentrations. The α-complementation assay (see Methods) was used to estimate the fidelity of the indicated RT with different concentrations of free Mg^2+^ (as indicated). The value for HIV-1 wild type (HXB2 clone) at 0.5 mM Mg^2+^ after background subtraction was set to 1 and higher values indicate a higher mutation frequency. Assays conducted with the RNA template, which go through 2 rounds of RT DNA synthesis, are shown on the left while those using the single round DNA templated assay are on the right (see Fig. 1). For wt HIV RT with the RNA templated system, the analysis was performed at several Mg^2+^ concentrations, while other RTs were with 0.5 and 6 or 0.5, 6, and 12 mM Mg^2+^ only (all free Mg^2+^ concentrations). Data used to construct this graph is shown in Table S1. * Relative mutation rate was based on the colony mutation frequency and was calculated from the proportion of bacterial colonies that were white or faint blue in the α-complementation assay divided by the total number of colonies (i.e., (white colonies + faint blue colonies) /(white colonies + faint blue colonies + blue colonies)). The background value (see Methods and Table S1) was subtracted from each condition to get the final value for comparisons. **Reverse transcriptases with amino acid changes are drug-resistant forms in an HXB2 background. AZTr: D67N/K70R/T215F/K219Q.

HIV and MuLV RT were also evaluated in the DNA templated assay. In this case, the mutation frequency difference was greater for HIV RT than in the RNA templated assay, with 6 mM Mg^2+^ producing an ~ 4-fold greater mutation frequency than 0.5 mM, while this difference was about 2-fold in the RNA assay. In contrast, and in agreement with the RNA templated system, MuLV RT values were essentially the same at both 0.5 and 6 mM Mg^2+^, as this enzyme demonstrated significantly higher fidelity than HIV RT only at 6 mM Mg^2+^. The greater difference at 0.5 vs. 6 mM Mg^2+^ in the DNA templated assay could result from differences in the fidelity of RT on DNA vs. RNA, however, they could also result from the starting RNA template having a higher mutation frequency than the starting DNA template (see Discussion).

In general, these results are in agreement with previous results from this lab [24] and others [25] demonstrating that the fidelity of HIV RT is greater in vitro when physiological free Mg^2+^ (i.e., ~ 0.5 mM, see Introduction) conditions are used. The results also support the notion that HIV RT demonstrates lower fidelity than MuLV and AMV RTs in vitro only when non-physiological high Mg^2+^ conditions are used [24].

### Analysis of mutations made by HIV RT in 0.5 and 6 mM Mg^2+^ using NGS

A SSCS NGS approach was used to determine the mutation profile of HIV RT wt using 0.5 or 6 mM Mg^2+^. This approach allowed the analysis of several thousand mutations from material analyzed in both the RNA and DNA templated assays described above. While we used an established approach that allows each RT synthesis event to be tracked using a random barcode [25], a new program to mine the data was generated (see Methods).

### Analysis of insertion/deletion (indel) mutations by NGS

The region analyzed for mutations is shown in Fig. 1. Nucleotides 6-160 of the sequence (155 total nucleotides) were evaluated for mutations. Six conditions (BKG control, and 0.5 or 6 mM Mg^2+^ for both the RNA and DNA templated system (Table 1)) were tested (see Methods for a description of how mutations were scored). This report emphasizes data from 1 experiment (Exp. 1), while a second set from an independent experiment (Exp. 2) yielded similar results (Table S2). The vast majority of mutations recovered by NGS were substitutions as opposed to indels (Table 1). We should note, however, that our protocol would likely eliminate large insertions and deletions as the material for analysis is excised from gels (see Methods). For this reason, only products that differed from the expected length by 5 nucleotides or less were evaluated in the NGS analysis, so even modestly large indels would have been excluded. As expected, indels occurred preferentially within runs of the same nucleotide on both the RNA and DNA templated samples and the positions mostly matched hotspots from previous mutational analysis with *lacZα* in cells [29]. Figure 3 shows the frequency of all substitution mutations for the RNA templated assay with 0.5 and 6 mM Mg^2+^ (excluding G>T and C>A mutations (see below)). Positions and mutations rates for the strongest (those in which at least one of the 2 conditions had a mutation frequency ≥1 x 10^−6^, these comprised great than 90% of all recovered insertions and about 60% of deletions) indel mutations are also shown (numbers on graphs placed above the specific base position with deletions underlined and insertions not underlined). Indel hotspots were almost exclusively in runs of the nucleotides, as expected. Most recovered insertions with both 0.5 (~ 91%) and 6 mM Mg^2+^ (~ 79%) occurred at positions 35-38 (insertion of A, 5′-GGG<u>AAAA</u>CCC-3′, 35-38 underlined) or 81-83 (insertion of T, 5′-ACCCCCC<u>TTT</u>CGCC-3′). Note that the exact position of the insertions within the run cannot be determined and the numbers in Fig. 3 are placed above the first base of the run. Deletions were more distributed with the 2 strongest sites being nucleotides 14-17 (deletion of T, 5′-CG<u>TTTT</u>AC-3′) and 75-80 (deletion of C, 5′-5′-A<u>CCCCCC</u>TTTCGCC-3′).

**Table 1.** Data from NGS Exp. 1.

| Condition<sup>a</sup> | Control DNA | DNA 0.5 mM Mg<sup>2+</sup> | DNA 6 mM Mg<sup>2+</sup> | Control RNA | RNA 0.5 mM Mg<sup>2+</sup> | RNA 6 mM Mg<sup>2+</sup> |
| --- | --- | --- | --- | --- | --- | --- |
| **Total Indices<sup>b</sup>** | 258,515 | 177,278 | 212,279 | 210,220 | 255,341 | 254,006 |
| **Total nucleotides<sup>c</sup>** | 4.007 x 10<sup>7</sup> | 2.748 x 10<sup>7</sup> | 3.290 x 10<sup>7</sup> | 6.517 x 10<sup>7</sup> | 7.916 x 10<sup>7</sup> | 7.874 x 10<sup>7</sup> |
| **Substitution Type** | **Number of recovered substitutions (substitution frequency per nucleotide x 10<sup>6</sup>)<sup>d</sup>** |  |  |  |  |  |
| A>C | 287 (7.163) | 581 (21.14) | 1227 (37.29) | 231 (3.545) | 1216 (15.36) | 2238 (28.42) |
| A>G | 332 (8.286) | 1500 (54.59) | 8835 (268.5) | 207 (3.176) | 4156 (52.50) | 10817 (137.4) |
| A>T | 134 (3.344) | 323 (11.75) | 1037 (31.52) | 41 (0.6291) | 1108 (14.00) | 2049 (26.02) |
| *C>A | 3459 (86.32) | 3310 (120.5) | 6051 (183.9) | 3597 (55.19) | 7043 (88.97) | 11804 (149.9) |
| C>G | 231 (5.765) | 1133 (41.23) | 8441 (256.6) | 226 (3.468) | 2040 (25.77) | 4427 (56.22) |
| C>T | 749 (18.69) | 1788 (65.07) | 12460 (378.7) | 540 (8.286) | 4796 (60.59) | 22619 (287.3) |
| G>A | 744 (18.57) | 5376 (195.6) | 8126 (247.0) | 371 (5.693) | 4231 (53.45) | 23202 (294.7) |
| G>C | 153 (3.818) | 253 (9.207) | 378 (11.49) | 54 (0.8286) | 386 (4.876) | 775 (9.843) |
| *G>T | 4773 (119.1) | 4212 (153.3) | 8069 (245.3) | 3245 (49.79) | 3274 (41.36) | 4261 (54.11) |
| T>A | 74 (1.847) | 280 (10.19) | 684 (20.79) | 50 (0.7672) | 1002 (12.66) | 1484 (18.85) |
| T>C | 321 (8.011) | 478 (17.39) | 1193 (36.26) | 181 (2.777) | 1760 (22.23) | 13537 (171.9) |
| T>G | 465 (11.60) | 623 (22.67) | 1564 (47.54) | 417 (6.399) | 1236 (15.61) | 3181 (40.40) |
| **Insertion Type** | **Number of recovered insertions (insertion frequency per nucleotide x 10<sup>6</sup>)<sup>d</sup>** |  |  |  |  |  |
| A | 0 | 16 (0.5822) | 47 (1.429) | 0 | 447 (5.647) | 500 (6.350) |
| C | 0 | 6 (0.2183) | 6 (0.1824) | 0 | 213 (2.691) | 333 (4.229) |
| G | 0 | 0 | 10 (0.3040) | 0 | 31 (0.3916) | 23 (0.2921) |
| T | 0 | 22 (0.8006) | 29 (0.8815) | 0 | 2692 (34.01) | 2605 (33.08) |
| **Deletion Type** | **Number of recovered deletions (deletion frequency per nucleotide x 10<sup>6</sup>)<sup>d</sup>** |  |  |  |  |  |
| A | 75 (1.872) | 86 (3.130) | 437 (13.28) | 27 (0.4143) | 273 (3.449) | 461 (5.855) |
| C | 37 (0.9234) | 49 (1.783) | 789 (23.98) | 23 (0.3529) | 745 (9.411) | 1082 (13.74) |
| G | 28 (0.6988) | 10 (0.3639) | 272 (8.267) | 23 (0.3529) | 201 (2.539) | 400 (5.080) |
| T | 74 (1.847) | 112 (4.076) | 485 (14.74) | 54 (0.8286) | 540 (6.822) | 1044 (13.26) |
| **Mutation Type** | **Percent of various types of mutations for each condition (mutation frequency x 10<sup>4</sup>)<sup>e</sup>** |  |  |  |  |  |
| *Substitutions |  | 98.4 (3.618) | 95.6 (12.49) |  | 79.3 (2.415) | 92.8 (10.35) |
| Insertions |  | 0.4 (0.0147) | 0.2 (0.0261) |  | 14 (0.4263) | 3.9 (0.4349) |
| Deletions |  | 1.2 (0.0440) | 4.2 (0.5485) |  | 6.7 (0.2040) | 3.2 (0.3568) |
| Overall mutation frequency |  | 3.677 x 10<sup>-4</sup> | 1.306 x 10<sup>-3</sup> |  | 3.045 x 10<sup>-4</sup> | 1.115 x 10<sup>-3</sup> |

a- Refer to Material and Methods for details. DNA assays included 1 round of RT DNA-template-directed DNA synthesis while RNA assays include an RT RNA-template-directed round and a second RT DNA-template-directed DNA synthesis round.
b- Indices are defined in Materials and Methods. To qualify as an “index”, the same 14-nucleotide barcode had to be recovered in 5 or more reads in NGS. Specific mutations were recorded when they appeared at a specific position in all the reads that constituted a particular index.
d- Insertion frequency per nucleotide was calculated by dividing the raw number of mutations of the specified type by the total number of nucleotides that were assessed for that condition. Unlike the raw number of mutations, this value allows a direct relative comparison of the data in the Table with higher numbers representing a greater frequency for that mutation. Note that the DNA and RNA templated assays cannot be directly compare with this number as the templates was synthesized by Q5 DNA polymerase or T3 RNA polymerase in the DNA and RNA assays, respectively. The Q5 synthesis product was used as the Control for both assays. Given the very low mutation rate of Q5, it is likely the RNA assays contain a significant proportion of mutations derived from T3 RNA polymerase that are not accounted for by the Control (see Results for further discussion).
e- e- Values were determined after subtracting background Control values from each mutation type using the frequency per nucleotide data above. These values were then summed for each of the 3 categories (i.e. substitutions (S), insertions (I), and deletions (D)). The percent values listed were calculated by dividing individual values by the sum of the 3 categories and multiplying by 100 (e.g. for Substitutions: (S/(S+I+D)) x 100). Mutation frequencies in parentheses were the individual S, I, and D values and the Overall Mutation Frequency was S+I+D for each category.
\*Due to high numbers in the Controls, C>A and G>T substitutions were excluded from data calculations (see Results).

**Figure 3.**
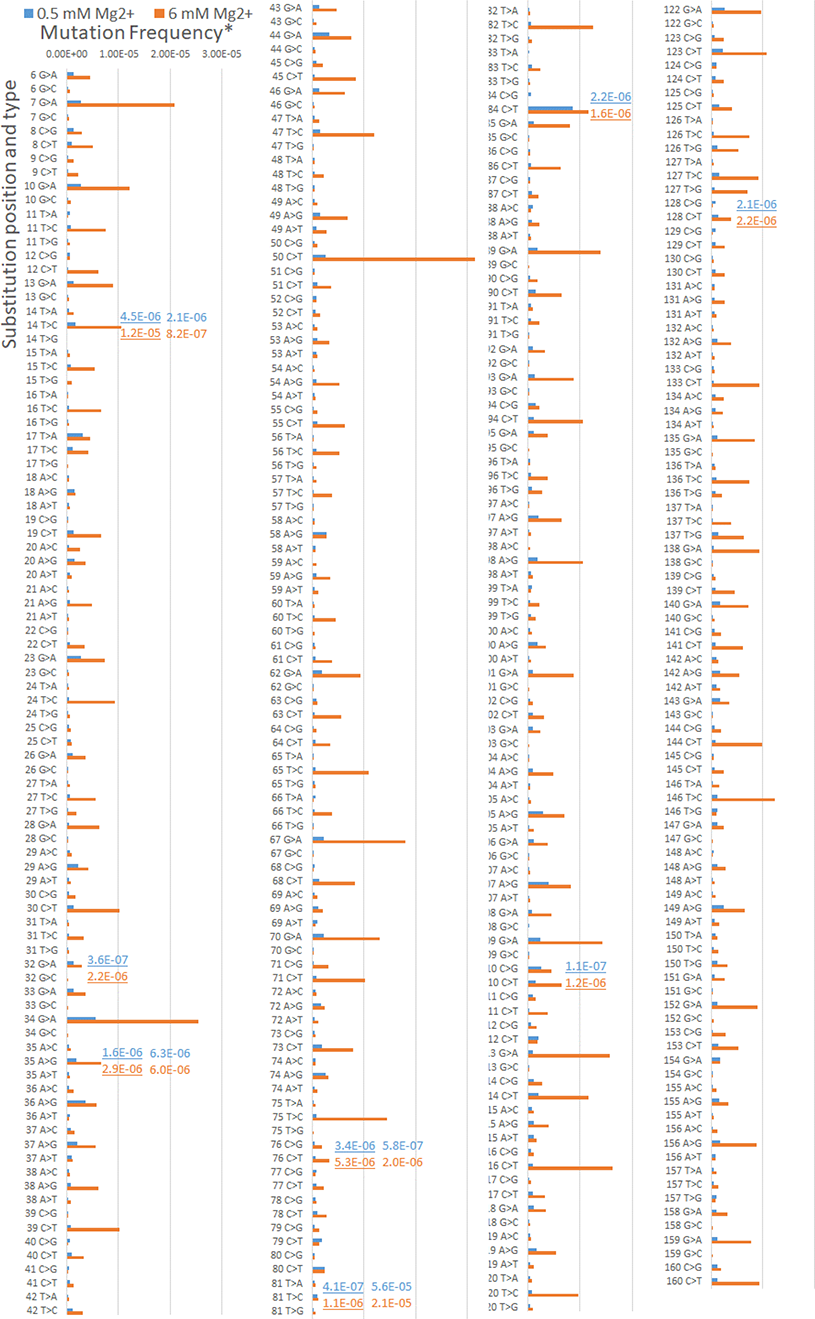
Mutation frequency with 0.5 or 6 mM Mg^2+^ in t e RNA templated assay. The mutation frequency per nucleotide at specific positions (see Table 1 and Fig. 1) for specific types of recovered substitution mutations at the two different Mg^2+^ concentrations (blue, 0.5 mM and orange 6 mM Mg^2+^) is plotted. Mutation frequencies for deletions (underlined) and insertions (not underlined) are given only for those positions where the rate at 0.5 or 6 mM was at least 1 x 10^−6^ for one or both of the conditions. Indel values were arbitrarily placed above the first nucleotide of a run of nucleotides although it is not known at which position the indel occurred. *Mutation Frequencies are for the specific mutation type and position and were calculated after subtraction of background rates at the same template position. See Table 1 for the total number of nucleotides scored for each condition.

For the RNA templated assay, the mutation frequency for indels did not change much between 0.5 and 6 mM Mg^2+^ while indels made up a greater percentage of total mutation in low Mg^2+^ due to the large increase in the substitution mutation rate in high Mg^2+^ (Table 1). Notable differences in the DNA templated assay were a much lower indel mutation frequency with low Mg^2+^, and a significant increase in deletions in high Mg^2+^. In Exp. 2 (Table S2), both insertions and deletions increased significantly at higher Mg^2+^ with DNA. Overall, in Exp. 1, indels comprised ~ 21% of all mutations at 0.5 mM Mg^2+^ in the RNA templated assay, but only 1.6% in the DNA assay under that condition. A similar trend was observed in Exp. 2 (Table S2). These results are consistent with the T3 RNA polymerase-derived RNA template containing a significantly higher level of indels than the Q5-derived DNA template [33] (see Discussion).

### Analysis of substitution mutations by NGS

Unlike indels, the substitution frequency increased ~ 4-fold in both RNA and DNA templated samples at 6 vs. 0.5 mM Mg^2+^ (Table 1). Background error frequencies, although relatively low for most mutation types, were much higher for C>A and G>T mutations. High C>A and G>T mutation rates are common in NGS analysis and are exacerbated by heating or other steps that damage or oxidize DNA [34–38]. Duplex-based NGS approaches as opposed to the SSCS approach used here, can more effectively correct for these errors, but these approaches are costly and can dramatically decrease coverage and yields [39, 40]. Because of the high background, G>T and C>A mutations were not included in calculations for mutation rates or proportions of the various mutation types shown in Tables 1 and S2, or Figs. 3 and 4. This is unlikely to significantly affect the mutation rate as G>T and C>A, like most other transversions, are low frequency mutations for RT and DNA polymerases in general.

**Figure 4.**
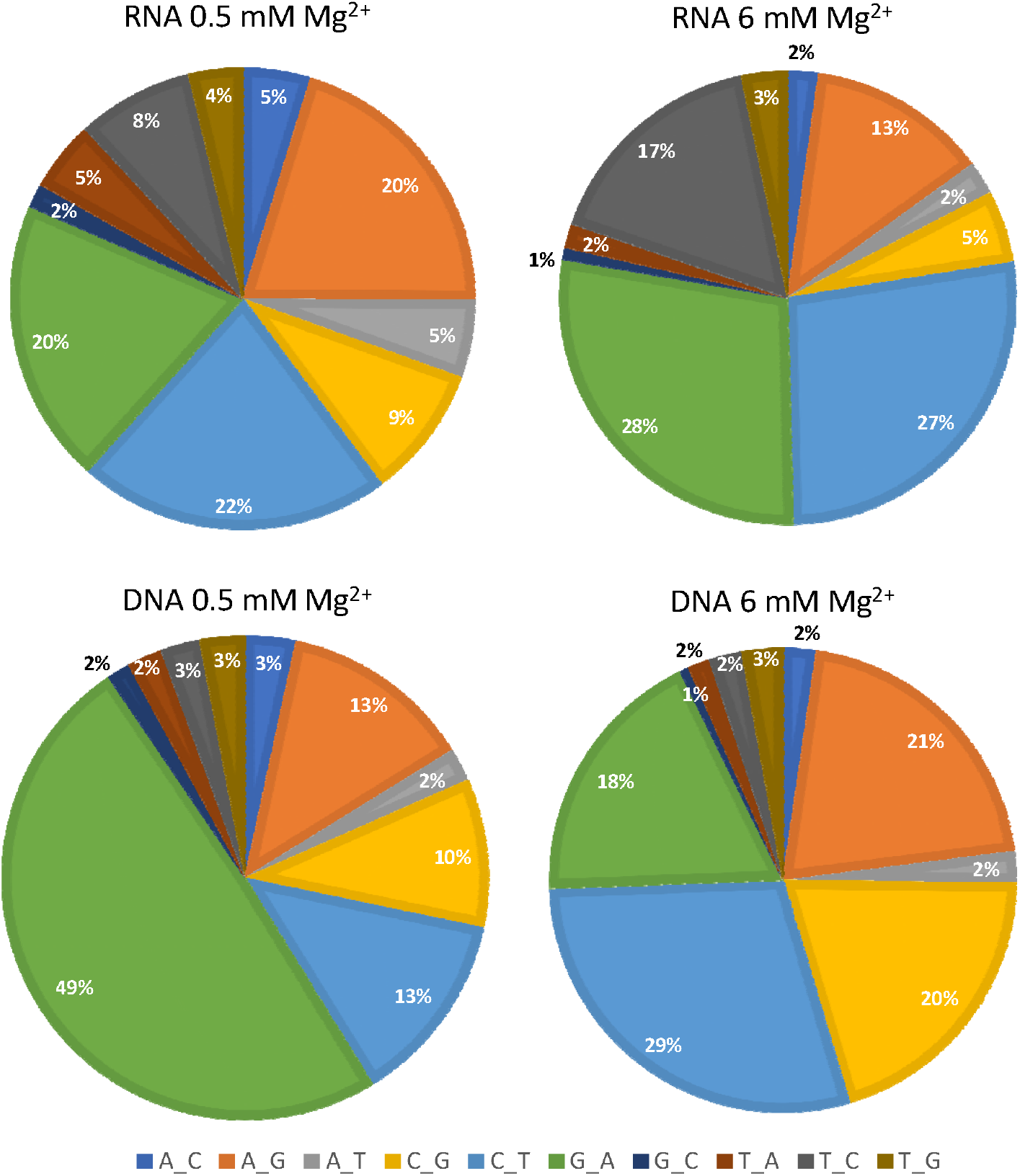
Proportion of various substitution mutations in the NGS assay. The proportions of substitution mutations recovered using the indicated conditions in the NGS assay are shown. G>T and C>A mutations are excluded from the analysis (see Results). See Table 1 for more information on the mutation rates for the various mutations.

As expected, transitions were more common than transversions for all conditions (Table 1 and Fig. 4 and Table S2 and Fig. S1 for Exp. 1 and Exp. 2, respectively). It is important to note that for the RNA templated assays, a particular mutation could have arisen during RNA or DNA-directed RT synthesis. For example, a G>A mutation could have resulted from a G coding for a T during RNA-directed DNA synthesis (the T would subsequently code for an A in the DNA-directed 2^nd^ round), or a C coding for an A during DNA-directed DNA synthesis (see Fig. 1). In contrast, RT-derived G>A mutations in the DNA templated assay arise from a G coding for a T during DNA-directed RT synthesis as this is the only RT round in that assay.

For the RNA templated assay, the relative proportion of various mutations was similar between the 0.5 and 6 mM Mg^2+^ conditions (Figs. 4). There was a modest increase in the relative proportion of G>A, C>T and T>C mutations with 6 mM Mg^2+^. As these transitions are among the most common mutation types for RT and many polymerases, it is possible that high Mg^2+^ has a greater effect on mutations that are easier to make.

For the DNA templated assay, the profile of mutation types showed a greater difference at 0.5 vs. 6 mM Mg^2+^ compared to the RNA templated assays (Fig. 4). The major difference between the results was the higher proportion of G>A mutations with 0.5 compared to 6 mM Mg^2+^. Also, there was a modest increase in A>G, C>T, and C>G mutations at 6 mM Mg^2+^, along with the corresponding decrease in the proportion of G>A mutations.

A comparison between mutation proportions at 6 mM Mg^2+^ between the RNA and DNA templated assay showed that it was similar, with the exception of T>C and C>G mutations (Fig. 4). The latter was relatively high in the DNA templated assays while the former was higher in the RNA templated assays. For the 0.5 mM Mg^2+^ assays, the most notable difference, once again, was the high proportion of G>A mutations in the DNA vs. RNA templated assay.

### Substitution mutation profile for the RNA templated assay at 0.5 and 6 mM Mg^2+^

An analysis of mutations with the RNA templated assays at individual nucleotides on the template suggested that, for many positions, the main effect of high Mg^2+^ was to magnify error-prone sites on the template rather than create new ones (Fig. 3 and see full data set for more information). DNA templated assays showed a similar trend (Fig. S2) although it was more difficult to assess, as the number of recovered mutations was not significantly above background at many sites (background was taken as the frequency of the same mutation type at the same position of the template in the control samples), especially for the 0.5 mM Mg^2+^ condition. An advantage of the RNA templated assay is that two rounds of RT synthesis are performed (Fig. 1), giving RT two chances to produce a mutation at each nucleotide position. Several of the strongest substitution mutations observed were C>T, A>G and G>A mutations, both in low and high Mg^2+^. These substitution types are common with HIV RT in vitro [41, 42] and cellular replication assays [28, 29, 43].

### Calculation of overall mutation rates in NGS assays for 0.5 and 6 mM Mg^2+^

The overall mutation frequencies and frequencies for particular types of mutations were calculated as described in the legend to Table 1. Overall mutation rates were modestly higher in the DNA templated assay, a trend that was also observed in Exp. 2 (Table S2). On a per nucleotide basis, in 0.5 mM Mg^2+^, HIV RT made approximately 1 mutation for every 2720 (1/(3.677 x 10^−4^)) and 3284 (1/(3.045 x 10^−4^)) nucleotides in the DNA and RNA templated assay, respectively. This rate rose to about 1 mutation for every 766 (1/(1.306 x 10^−3^)) and 897 (1/(1.115 x 10^−3^)) nucleotides in 6 mM Mg^2+^ for DNA and RNA template assays, respectively. These rates would exclude G>T and C>A mutations as well as larger insertions/deletions that cannot be tested by this approach (see Methods). In all cases, indel mutation rates were relatively low, although they made up ~ 21% of total mutations (after BKG subtraction) in the 0.5 mM RNA templated assay. Exp. 2 demonstrated a similar trend although mutation rates in DNA assays were higher than those in RNA assays to a greater extent than in Exp. 1 (Table S2).

### Comparison of the NGS results with other results in cellular assays

In general, the mutation rate of HIV is much lower when measured during replication of HIV in cells than using HIV RT in vitro. The sensitivity of HIV RT to Mg^2+^ concentration and the fact that most in vitro assays have been done in high Mg^2+^ is likely part of the reason for this (see Introduction). In contrast to in vitro assays, most cell culture experiments have identified a relatively small number of mutations, however, more recent SSCS NGS cellular approaches identified larger mutation sets [43], while an extensive analysis of mutation types using *lacZα* complementation in cells was also performed by Abram et al. [29]. Our experiments used a region of *lacZα* nearly identical to Abram et al. (Fig. 1), allowing a direct comparison of the results. The numbering in Fig. 1 is consistent with numbering in Abram et al. (see Methods). Abram et al. [29] recovered a total of ~ 600 substitution mutations (including those recovered with *lacZα* in the forward and reverse orientations) at 71 of the 174 nucleotide *lacZα* region in their assay (“single nucleotide substitutions” or “class 1” mutations in that report), and indels at a much lower rate (~ 5% of recovered mutations, “class 2” mutations). Since the assay was based on α*-*complementation, only nucleotide changes resulting in detectable changes in *lacZα* activity were recoverable, unless the mutation was coupled with another detectable mutation as part of a *lacZα* gene with multiple mutations. Although Abram et al. [29] also recovered several mutations as part of multiple nucleotide substitutions in the same *lacZα* gene (“class 3” mutations in that report), we choose to compare our NGS mutations to those recovered as single nucleotide substitutions, or single nucleotide frameshifts (class 1 and 2 mutations, respectively), which constituted the bulk of indel mutations recovered by Abram et al. [29].

There was a strong correlation for signal nucleotide insertion mutations with the two predominant hotspots from Abram et al. [29] constituting a large proportion of the indels recovered with NGS (nts 35-38 and 81-83 discussed above). Unlike Abram et al. [29] where deletions were fewer compared to insertions, our NGS assay recovered a similar level of insertions and deletions (Tables 1 and S2). Perhaps most notable, was the low indel mutation rates in both our assay and the Abrams et al. [29] assay. Only the 0.5 mM RNA templated assays showed indel mutations rates greater than ~ 8% of total mutations in the NGS assays, comparable to the ~ 5% for “class 2” mutations in Abrams et al. [29]. Many in vitro assays using *lacZα* demonstrate higher indel rates [41, 44, 45].

Substitution mutations were more difficult to correlate as our NGS assay does not depend on *lacZα* activity, essentially detecting all mutations. But since the assay in Abrams et al. [29] was based on α*-*complementation, only nucleotide changes resulting in detectable changes in LacZα activity were recovered, unless the mutation was coupled with another detectable mutation as part of a *lacZα* gene with multiple mutations. Unlike substitutions, indels, especially those closer to the start of the *lacZα* gene, always inactivate the gene leading to a white colony in the α*-*complementation assay. We took the approach of comparing only the highest frequency mutations from NGS with those from Abram et al. [29]. In the NGS assay for the RNA templated sample with 0.5 mM Mg^2+^ (this condition was used as it most closely matches cellular replication), a total of 332 different types of mutations were present above background at 155 total positions, with at least 1 mutation type recovered at each position that was scored. For “high-frequency mutations”, an arbitrary mutation frequency cut-off for specific mutation types at a specific nucleotide position of 1.25 x 10^−6^ was used, which constituted the 61 highest frequency substitution (about 18% (61/332) of all the types of recovered mutations) in the NGS assay (Fig. 3). These were compared to substitution mutations from Abram et al. [29] using the following criteria: only mutations recovered at least 1 time in the Abram et al. [29] α*-* complementation assay were classified as “detectable” in that assay. Of the 61 highest frequency mutations in the NGS assay, 39 were not detected in the Abrams et al. [29] assay, and therefore could not be compared between the two assays. Of the remaining twenty-two, 7, 6, 3, and 6, were recovered 1-2, 3-5, 6-11, and 11 or more times, respectively, in the Abram et al. [29] assay. For perspective, there were 33 mutations (of the ~ 120 different mutation types recovered as class 1 mutations) in the Abram et al. [29] assay that were recovered 6 or more times and 16 recovered 11 or more times.

## Discussion

This report confirms and expands upon previous results showing that HIV RT, but not MuLV and AMV RTs, demonstrate greater fidelity in Mg^2+^ concentrations that more closely match the free Mg^2+^ concentrations in cells. Common drug resistance mutations in HIV RT demonstrated a similar behavior as did HIV-1 subtype A/E RT, HIV-2 RT and PFV RT, although the latter showed statistically higher fidelity than the other RTs used in this report (Table S1). All HIV-1 drug resistance mutants were less accurate in higher Mg^2+^ and, consistent with previous results [31], only K65R RT demonstrated a statistically lower mutations rate than wt HIV-1 RT (Table S1). Although some drug resistant mutants (M184V in particular), showed a greater decrease in fidelity than wt in high Mg^2+^, none of the mutants were dramatically more affected by Mg^2+^ than wt. We have previously shown that the potency of NRTIs diminished in 0.5 mM Mg^2+^ compared to 6 mM [23], while the opposite occurs for NNRTIs [22, 23]. This suggests that interactions involving Mg^2+^ are important for modulating the potency of these drugs, and advocates for the possibility of resistance mutations arising to exploit these properties to further diminish drug potency. Although the experiments here did not directly test this, they do show that the tested mutants retain the pronounced shift to lower fidelity at higher Mg^2+^ concentration. Therefore, none of the mutants tested here suggest an altered interaction with Mg^2+^ relative to wt, at least in the context of fidelity.

We previously postulated that the enhanced fidelity of HIV RT at lower Mg^2+^ could result from the lower velocity of the reaction, which increases the residence time on each template base and therefore affords better discrimination [24]. More recent results indicate that the putative A (catalytic Mg^+2^) and B (nucleotide bound Mg^2+^) divalent cation binding sites in the polymerase domain of HIV-1 RT have dramatically different affinities for Mg^2+^ [46], with sites A and B binding with K_d_s of 3.7 mM and 29 μM, respectively. The authors propose that “weak binding of the catalytic Mg^2+^ contributes to fidelity by sampling the correctly aligned substrate without perturbing the equilibrium for nucleotide binding at physiological Mg^2+^ concentrations”. These findings are consistent with our previous hypothesis. Further, the range of Mg^2+^ concentrations explored here (from 0.25 to 12 mM) for wt HIV RT (Fig. 2) and in our previous work [24] span the proposed K_d_ for site A. This mechanism is also consistent with there being little change in fidelity between 6 mM and 12 mM Mg^2+^ for HIV-1 RT (Fig. 2) as the occupancy of site A would likely change little over this range, assuming the K_d_ value for this site is close to the predicted value of 3.7 mM. Since MuLV and AMV RTs do not demonstrate large fidelity changes between 0.5 mM and 6 mM Mg^2+^ (Fig. 2), this may suggest that the binding affinity for Mg^2+^ at sites A and B differ for these enzymes compared to HIV RT, but this remains to be determined.

The mutation frequencies in the α-complementation assay were similar for most of the enzymes tested. Only PFV RT at 0.5 mM and 6 mM Mg^2+^, K65R at 6 mM Mg^2+^, and MuLV RT at 6 mM Mg^2+^ (in both the RNA and DNA templated assays) showed statistically greater fidelity than HIV-1 RT wt at the same Mg^2+^ concentration, while M184V RT was statistically less accurate at 6 mM Mg^2+^ (Table S1). The α-complementation assay examined mutations over a modestly larger sequence range than the NGS assay (Fig. 1). Ignoring this small difference and using the same 155 nucleotide scored range (see Results), the overall mutation frequency for HIV-1 wt on the RNA templated system after background subtraction for 0.5 mM and 6 mM would be approximately 1.2 x 10^−5^ and 2.5 x 10^−5^ ((colony mutation frequency-BKG)/310 nucleotides (2 rounds of RT synthesis over the scored region), respectively. In the DNA templated assay these values would be 0.64 x 10^−5^ and 2.6 x 10^−5^ (note there is only a single round of RT synthesis in this assay so the total nucleotides would be 155), respectively. These number are likely to be significantly lower than the real mutation rate as most mutations made by RT are not detected in the α-complementation assay (see Results). The numbers are in the same range as number we previously calculated using a similar assay [24]. Interestingly, these mutations rates are approximately the same as those calculated for HIV infection in cell culture by Abram et al. [29] using a α-complementation assay.

The NGS analysis allowed for a direct comparison of the frequency and types of mutations made in both RNA and DNA templated assays with 0.5 mM or 6 mM Mg^2+^. Only wt HIV-1 RT was examined with NGS so it is not possible to determine if RT drug resistance mutations affected the types of mutations that were made. An interesting finding was that unlike substitutions which increases about 4-fold overall, there was no significant increase in indel mutations between 0.5 mM and 6 mM Mg^2+^ in the RNA templated assay, and indels made up a lower proportion of the total mutations at 6 mM Mg^2+^, owing to that condition having more substitutions (Table 1). In contrast, indels increased significantly at the higher Mg^2+^ concentration in the DNA templated assay. There was more mutation data for the RNA vs. DNA templated assay, probably owing to two vs. one round of RT synthesis in this assay. Also, the data between the two experiments (see Table S2 for second exp.) matched better with the RNA, especially for indel mutations. These discrepancies with the DNA data lend some uncertainty to the results. However, one thing that was especially clear was the much lower indel rate with DNA vs. RNA at 0.5 mM Mg^2+^ (Tables 1 and S2). This may be due to the different mechanism for indel vs. substitution mutations. The former result mainly from slippage of the template or primer strand while the latter can result from slippage and other mechanism [47, 48]. Indels typically occur by addition of the “correct” nucleotide on a transiently misaligned primer-template, followed by continued elongation prior to realignment. Substitutions can occur by this mechanism when realignment occurs prior to elongation, or by direct incorporation of an incorrect nucleotide. Indels could be more pronounced on RNA, especially if slippage occurs more frequently. An alternative explanation, and one we favor, is that the RNA template in the RNA assays may have more indels (and substitutions) to start, due to the lower fidelity of T3 RNA polymerase used to produce the RNA template vs. Q5 DNA polymerase used to produce the DNA template. The substitution mutation rate of Q5 DNA polymerase has been reported to be ~ 5 x 10^−7^, lower than any other thermostable polymerase [30]. Q5 contributions to errors in the DNA templated assays should be very small as it has ~ 2 orders of magnitude greater fidelity than HIV RT. As noted in the Results, there are many steps in the NGS assay and manipulations such as heating and running material on gels as well as nucleic acid oxidation and PCR cycling could contribute to errors observed in the control assays (see [30] for a detailed analysis of error sources in PCR assays), beyond errors that were produced by Q5 which we expect would be very low. Although the fidelity of T3 and other RNA polymerase has not been directly measured (this is difficult because of the necessity to reverse transcribe the RNA to DNA for sequence analysis), recent experiments suggest phage RNA polymerases contribute a significant level of mutations to reverse transcriptase fidelity assays that use phage polymerase-derived RNAs as the template [33]. The mutation rate of human RNA polymerases II is also unknown and its contribution to HIV’s mutation spectrum remains unclear. Note that the above discussion would also predict that the mutation rate in the RNA templated assay should be higher than the DNA templated assay since the starting template had more mutations. Despite this, in the NGS experiments, DNA templated assays showed small, but consistently higher mutation rates (Tables 1 and S2). This was not observed in the α-complementation assay, but those results were from a much smaller data set. One possibility is that HIV RT is less accurate when copying DNA than RNA, as has been suggested previously [33, 49]. Since the numbers in our assays were not dramatically different with RNA and DNA, more data and a better understanding of the mutation rate of phage RNA polymerase would be required to address this question.

With respect to the proportion and positions of indel mutations, the NGS results compared reasonably well with results from Abram et al. using the same sequence in HIV cell replication assays [29]. Unlike some previous in vitro results with HIV RT [41, 44, 45], indels made up a relatively low proportion of the observed mutation in our NGS results (Table 1 and S2), as they did in Abram et al. [29]. Notably, other in vitro NGS results with HIV RT have yielded a low proportion of indels as well [42]. A possible explanation for previous high indel rates in vitro is that most of those assays were performed using α-complementation and indels are easier to detect in those assays (see Results), although this is not consistent with the low indel rates observed by Abram et al. [29]. Perhaps the subjective nature of the α-complementation assay can lead to variation in the results, especially if all the colonies scored as mutations are not sequenced.

Substitution mutations were more difficult to compare between our NGS assay and other cellular assays. Like other NGS assays which use HIV-1 RT in vitro [25, 42], as well as other cellular replication assays [28, 29, 43], our NGS results showed a strong tendency for transitions vs. transversions, and a high proportion of G>A and C>T mutations (Fig. 4). However, comparisons to the Abram et al. [29] results indicated that the majority (39 of 61 (see Results)) of the high frequency mutations (arbitrarily set as those with mutation frequencies ≥ 1.25 x 10^−6^) observed in the RNA templated assay with 0.5 mM Mg^2+^ were not found by Abram et al. using α-complementation and, therefore, could not be compared. Nine of the remaining 22 mutations from that group were found 6 or more times in the Abram et al. [29] results while the other 13 were recovered less than 6 times. A complete correlation would not be expected even if our in vitro conditions perfectly mimicked cellular conditions as there are factors in the cell (discussed in Abram et al. [29]) that are likely to modify the mutation spectrum. It would be interesting to use NGS over the same sequence to directly compare both in vitro and cellular results. This would allow a comparison of thousands of mutations and allow a more direct analysis of the contribution of RT vs. cellular factors in the mutation spectrum of HIV.

The mutation rate observed with NGS was also somewhat higher than expected based on results with α-complementation. The estimated rate of 1.2 x 10^−5^ in the α-complementation in the 0.5 mM Mg^2+^ RNA templated assay would necessarily underestimate the actual mutation rate as this assay only scores a fraction of the substitution mutations (see Results). However, the rate for the same condition in the NGS assay which detects “all” mutations, was ~ 20-fold higher (3.045 x 10^−4^ and 1.827 x 10^−4^, Exp. 1 and 2, respectively (Table 1 and S2)). Abram et al. [29] recovered about 110 different types of substitution mutation (class 1 mutations in their data) over the same range we used in our NGS assay. Presumably our assay could have recovered 465 (155 x 3) different types of substitutions over that range. Using this logic, which does not take into account the mutation frequency at specific nucleotide locations, the NGS assay should detect roughly 3 times more substitutions. This number is likely to be higher as nearly all detected mutations were in the first two-thirds of the scored region in the Abram et al. [29] assay, suggesting that α-complementation cannot detect many mutations near the *lacZ* C-terminus. Still, a 5-fold increase in NGS would seem more likely than the observed 20-fold. The overall mutation frequency from our results using 0.5 mM Mg^2+^ compared well to other NGS results with HIV RT [25], but those were conducted with higher Mg^2+^ concentrations, while NGS results with lower Mg^2+^ produced a lower mutation rate than we found [25, 42]. An NGS analysis of HIV replication in cells over a region of the HIV integrase gene has also been conducted using a SSCS approach similar to our NGS approach to correct for errors derived from amplification and sequencing [43]. Only transition mutations were recovered at high enough frequencies above background to analyze. The transition mutation rate for NL4-3 HIV-1 was ~ 1 x 10^−4^, while other HIV-1 subtypes showed several-fold lower mutation rates. Since this is the rate for transitions only (which likely constitute a large portion of the total mutations), the rate for all mutations would be higher. It would be interesting to test the RTs from some of the other subtypes to see if they also demonstrate higher fidelity in vitro. Again, an analysis of the same gene region in vitro and in cellular replication using NGS approaches that decrease background mutations like the SSCS used here or duplex sequencing analysis [39, 40] would be very helpful for these comparisons.

## Materials and Methods

### Materials

Deoxyribonucleotide triphosphates (dNTPs) were from Roche. Q5 DNA polymerase, T4 polynucleotide kinase (PNK) calf intestinal phosphatase (CIP), restriction enzymes NlaI, High Fidelity PvuII, EcoRI, and EcoRV, NEBNext Ultra II DNA Library Prep Kit for Illumina (Cat# E7645S), and NEBNext Multiplex Oligos for Illumina (Cat# E7335S) kit were from New England BioLabs. RNase-DNase free was from MilliporeSigma. SYBR Safe DNA gel stain was from Invitrogen. Radiolabeled ATP (γ-^32^P) was from Perkin-Elmer. G-25 spin columns were from Harvard Apparatus. The RNA purification kit was from GeneJET. All DNA oligonucleotides were from Integrated DNA Technologies (IDT). Wild type HIV-1 reverse transcriptase (RT) (from HXB2 strain) was prepared as described [50]. Aliquots of HIV RT were stored frozen at −80°C and fresh aliquots were used for each experiment. Prototype foamy virus reverse transcriptase was kindly provided by Dr. Eddy Arnold (Rutgers University). All other enzymes were obtained as clones and kindly provided by Dr. Stefan Sarafianos (Emory University), or Dr. Stephen Hughes (National Institutes of Health). They were purified and stored using methods similar to wt RT. All other chemicals were from VWR™, ThermoFisher Scientific, or MilliporeSigma.

### Construction of plasmid pBS∇EcoRV_567_ for *lacZ*α complementation assays

Plasmid pBS∇EcoRV_567_ was derived from pBSΔPVUII [51] by removing a portion of the plasmid between the lone remaining PVUII site (position 764 of the pBSM13+ parent plasmid) and the single NaeI site at position 564. These cleavages leave blunt ends and an insert was used to replace the removed segment. The insert was made using primers 5□-CTGGCGTAATGCGAAGAGG-3□ and 5□ - <u>GATATCTTATTA</u>GCGCCATTCGCCATTGAGGC-3□ and performing PCR (using Q5 DNA polymerase and manufactures protocol) using the pBSΔPVUII plasmid as template. The underlined region of the second primer coded for the new EcoRV site and two consecutive stop codons to terminate *lacZ*α synthesis. Plasmid constructs were grown in GC5 bacteria and individual clones were sequenced to obtain plasmids with the insert in the correct orientation. The resulting pBS∇EcoRV_567_ plasmid (see Fig. 1) retains the multiple cloning site (MCS) of pBSM13+ but the *lacZα* gene segment now mimics the gene in plasmid pNLZeoIN-R-E-.LZF/R used by others to analyze HIV RT fidelity in cell culture [29]. Nucleotides 3-162 of *lacZα* (forward) from pNLZeoIN-R-E-.LZF/R (numbering is as described in [29] where “1” designates the start of the first *lacZα* codon (6^th^ actual codon)) after the multiple cloning site (MCS)) are the same in pBS∇EcoRV_567_ while nucleotides 163-174 are not present in pBS∇EcoRV_567_. Addition amino acid coding nucleotides upstream of nucleotides 3-162 (including the start of the *lacZα* gene and the MCS) are not identical to those in pNLZeoIN-R-E-.LZF/R. These differences had no noticeable effect on α-complementation.

### End-labeling of oligonucleotides with T4 PNK

DNA oligonucleotides were 5′ end-labeled in a 50 μl volume containing 50-100 pmol of the oligonucleotide of interest, 1× T4 PNK reaction buffer (provided by manufacturer), 10 U of T4 PNK and 10 μl of (γ-^32^P) ATP (3000 Ci/mmol, 10 μCi/μl). The labeling reaction was done at 37°C for 30 min according to manufacturer’s protocol. PNK enzyme was heat inactivated by incubating the reaction at 75°C for 15 min. Excess radiolabeled nucleotides were then removed by centrifugation using a Sephadex G-25 column.

### Production of the RNA template for *lacZα* complementation assay

Plasmid pBS∇EcoRV_567_ was cleaved with NdeI and T3 RNA polymerase was used to make run-off transcripts ~ 644 nucleotides in length using the manufacturer’s protocol. RNA was isolated using an RNA purification kit and quantified by UV absorption. The integrity and size of the RNA was confirmed by denaturing gel electrophoresis.

### Two round *lacZα* complementation assay (RNA templated system)

This assay measures 2 round of DNA synthesis by RT. Conditions for extension with RT were: 50 mM Tris-HCl pH 8 (final pH in reactions at 37°C was ~ 7.7), 80 mM KCl, 1 mM DTT, 20 μM dNTPs, 0.5 or 6 mM MgCl_2_ (final free concentration in reactions after correcting for chelation by dNTPs [23]), 37°C for 45 minutes. Round 1 used the RNA template described above to make the first strand of DNA. Four identical reactions were carried out for each condition using 25 nM template and 50 nM primer (round 1 primer below 5′-^32^P-labeled at low specific activity). Primer and template were hybridized in reaction buffer without dNTPs and MgCl_2_ by heating to 65°C for 5 min then slow cooling to room temperature in a heat block. Reactions were initiated by addition of RT (final concentration 100 nM). One μg of RNase A was then added to each reaction and incubated for 5 min then heated to 65°C for 2 minutes. After addition of 2X loading buffer (90% formamide, 10 mM EDTA (pH 8), 0.025% bromophenol blue and xylene cyanol) the samples were run on a 6% denaturing polyacrylamide gel [52]. Fully extended DNA (289 nucleotides) was located with a phosphorimager and excised then isolated by the crush and soak method [52]. The DNA was then used as a template to make a second strand of DNA (252 nucleotides) using the buffer conditions described above and the round 2 primer shown below. The round 2 primer was 5□ −^32^P end-labeled at ~ 10-fold greater specific activity than the round 1 primer to make it easier to locate and separate from the 289-nucleotide template. Two pmoles of round 2 primer was prehybridized (10 μl of 50 mM Tris-HCl pH=8, 80 mM KCl, 1 mM DTT, 80°C, 3 min, then decrease temperature by 2°C/minute to 37°C in a PCR machine) to the recovered DNA from the 4 round 1 reactions and round 2 reactions were carried out as described above. The second round DNA product was separated from the round 1 template using an 8% denaturing polyacrylamide gel and recovered as described above. The recovered DNA was quantified based on radioactivity and PCR amplified using high fidelity Q5 DNA polymerase. The PCR products were processed by cleavage with EcoRI and EcoRV and purified on native gels. Plasmid pBS∇EcoRV_567_ was cleaved with the same restriction enzymes and dephosphorylated with PNK. The purified PCR-derived inserts were then ligated into the cleaved plasmid and used to transform competent E. coli capable of α-complementation. Standard blue-white screening was used to score mutations. Details for these steps have been published [24]. The main differences are that plasmid pBS∇EcoRV_567_ was used here instead of pBSΔPVUII, the insert to vector ratio was 3:1, and restriction enzymes were “High Fidelity” versions of EcoRI and EcoRV while EcoRI and PvuII were used in the previous system. The different restriction enzymes and plasmid sequences required the use of different 5□-^32^P end-labeled primers to prime the 2 rounds of RT DNA synthesis: round 1: 5□-GATTTAGAGCTTGACGGGGA-3□, producing a full-length extension product of 289 nucleotides; round 2: 5□-AGGATCCCCGGGTACCGAGC-3□, producing a full-length extension product of 252 nucleotides. The difference in product lengths was used to separate the products using denaturing gel electrophoresis. These same primers were used to produce the PCR products described above. Bacterial colonies were scored by blue-white screening as previously described [24]. In some cases, the results (Table S1) were from a single experiment while in others they are summed values from more than one independent experiment. Statistical analysis to assess significance was conduct using a Chi-squared tests.

### Production of single stranded DNA for single round *lacZα* complementation assay and single strand consensus sequencing (SSCS) Next Generation Sequencing (NGS)

Single stranded DNA was produced using Q5 DNA polymerase and the manufacture’s recommended protocol. First, a 306 base pair dsDNA product was made by PCR using the primer from round 1 above and a second primer: 5’- AATTAACCCTCACTAAAGGG. Plasmid pBS∇EcoRV_567_ (0.1 μg) was used as template. Reactions were in 50 μl with 1 unit of Q5 DNA polymerase. Cycles were 94°C, 5 min, then 15 cycles of 94°C, 55°C, and 72°C for 30 seconds each, followed by 1 cycle of 72°C for 5 min. Cycling was minimized in order to decrease PCR-derived mutations. Products were run on an 8% non-denaturing polyacrylamide gel [52] using SYBR Safe DNA gel stain which allowed visualization of the products using blue LED light. Excised products were recovered using the crush and soak method [52] and quantified by UV absorbance. One pmole of dsDNA product was used in an asymmetric PCR reaction with 50 pmoles of 5′-^32^P end-labeled (low specific activity) round 1 primer above. Conditions were as above but 30 cycles were used. The reactions (typically 3 reactions were combined) were phenol extracted and precipitated with ethanol using standard procedures. The recovered material was digested with EcoRI (20 unit) for 20 min at 37°C, then run on a 6% denaturing polyacrylamide gel along with a size marker to locate the correctly sized DNA (using a phosphorimager) which was excised and recovered as described above. The digestion with EcoRI was used to cleave any double stranded DNA so as not to contaminate the desired 306 nucleotide single stranded product with complementary DNA that could interfere with downstream applications. The ssDNA product was quantified by specific activity.

### Single round *lacZα*-complementation assay (DNA templated system)

The assay was carried out using the round 2 conditions for the “two round *lacZ*α complementation assay” above. It essentially mimicked the second round of DNA synthesis except that the starting template was the single stranded DNA produced by asymmetric PCR as described above (only a single 25 μl reaction was typically performed for each condition). This DNA was primed with the 5□-^32^P end-labeled round 2 primer and the 252 nucleotide fully extended ssDNA product was isolated and processed as described above.

### Sources of mutations in the *α*-complementation assays

Two different systems were used to test fidelity (see Methods and Fig. 1). The first (termed “RNA templated” assay) was used for all RTs that were tested and started with an RNA template produced with T3 RNA polymerase from pBS∇EcoRV_567_. This template was copied with RT to produce a DNA that, after isolation, was used as a template to make a second DNA strand. The second strand was used as a template in a PCR reaction with Q5 DNA polymerase. The resulting purified product was processed as described above and used as a plasmid insert for the α*-*complementation assay. The second (termed “DNA templated” assay) system started with a DNA template produced by asymmetric PCR with Q5 DNA polymerase. This DNA was used as a template for RT to produce a second DNA that was then used for PCR as described above. The first system essentially mimics the entire reverse transcription process while the second mimics 2^nd^ strand synthesis only. With respect to background mutations, the starting templates in each system would include mutations present in plasmids which are likely insignificant. The first system also includes mutations made by T3 RNA polymerase while the second has mutations made during asymmetric PCR with Q5 polymerase. Both systems would include background generated by the PCR steps as well as those related to the assay steps (e.g., restriction digest and ligation). The background controls (see Table S1) for each assay would correct for all mutations occurring after those present in the starting material. In the second system, the mutation rate in the starting DNA template is also accounted for as the PCR step uses that template to produce the control insert. However, errors made by T3 RNA polymerase are not accounted for in the first system. Importantly, recent results suggest that phage RNA polymerases contribute significantly to mutations in fidelity assays using RNA as a template [33]. Results in Table S1 with 0.5 mM Mg^2+^ and HIV RT wt showed colony mutation rates of 0.00435 and 0.00189 for the RNA and DNA template systems, respectively. However, it is not possible to directly assess what proportion of the higher error rate in the RNA system was due to errors in the original RNA template vs. those contributed by the extra RT-directed RNA to DNA synthesis step in the RNA system. Overall, we expect that the real background in the RNA templated system would be significantly higher than the DNA templated system and is higher than the 0.00087 frequency for the background control in Table S1. Therefore, the mutation frequencies for this RNA system in Table S1 (and Fig. 2), which subtracted the control background, would represent a maximum frequency. They are probably modestly high for the 6 mM Mg^2+^ conditions, which had ~ 10-fold higher mutation frequencies than the background control, and more significantly high for the 0.5 mM Mg^2+^ conditions, which were only ~ 4-fold above background.

### Production of RT DNA products for NGS

Assay conditions for producing the DNA products for NGS were the same as those described above except that the 5′-^32^P end-labeled primer 5’-AAAAGGTAGTGCTGAATTCGATCACGNNNNNNNNNNNNNNGTGAGTCGTATTACAA TTCA-3′ (14N barcode primer) was used to primer the second synthesis round in the two round assay and the only synthesis round in the single round assay. This primer contains a 14-nucleotide random barcode region that is used for tracking individual NGS products for fidelity analysis (see below and [25]). This corresponds to 4^14^≍2.7×10^8^ different potential barcodes for RT synthesis products. The 3′ terminal nucleotide of the primer is 1 base upstream of *lac*Zα position “1” in pNLZeoIN-R-E-.LZF/R, allowing a direct comparison of these results with results in cell culture assays using that plasmid [29].

### Production of ssDNA control for NGS

The same control was used for the single and two round *lacZα* assays to set the background mutation rate in the NGS analysis. One pmole of the ssDNA made by asymmetric PCR (see above) was used as a template for a single round of Q5 DNA polymerase extension using 2 pmoles of 14N NGS index primer from above. One cycle of PCR (95, 50, and 72°C, for 1 min) with 1 unit of Q5 DNA polymerase was performed in 50 μl of Q5 supplied buffer and the correct length extension product was isolated from a 6% denaturing polyacrylamide gel as described above. Note that for the two round assay, this control does not include any mutations generated during the production of the RNA template that was used at the start of the assay (see above). Therefore, background errors from the RNA template are not subtracted away in the assay results.

### Production of PCR products and addition of adapters for NGS

For NGS, the final DNA products from the single or two round *lacZ*α complementation assays described under “Production of RT DNA products for NGS” were used to produce PCR products with the following primers: forward, 5’-GACGGGGAAAGCCGATATCTTATTA-3’ and reverse, 5′-^32^P end-labeled 5′-AAAAGGTAGTGCTGAATTCG-3′. The 3′ nucleotide of the forward primer is one base downstream of nucleotide 162 in pNLZeoIN-R-E-.LZF/R. Therefore, the PCR products produced contain nucleotides 1-162 of the *lacZ*α gene from pNLZeoIN-R-E-.LZF/R and, in our system, these nucleotides were derived from one or two rounds of synthesis with RT. Conditions for PCR reactions with Q5 DNA polymerase were 94°C, 5 min, then 94°C, 55°C, and 72°C for 30 seconds each, followed by 1 cycle of 72°C for 2 minutes. Reactions were conducted for 20, 22 or 24 cycles with aliquots being removed after the 72°C for 2 minutes extension step. Fifty pmoles of each primer were used in a 50 μl final volume and duplicate PCR reactions were performed for each condition in order to get a higher yield of products. For all conditions, 1.5 x 10^6^ (calculated from specific activity of the radiolabeled primer) molecules of template DNA from RT reactions was used in each 50 μl PCR reaction. PCR products (244 nucleotides) were separated on an 8% nondenaturing polyacrylamide gel. PCR products of the correct size were typically excised from either the 20, 22, or 24 cycle lanes, depending on which yielded the most material. Recovered products were processed using the “NEBNext Ultra II DNA Library Prep Kit for Illumina (Cat# E7645S)” while adapters used were from the “NEBNext Multiplex Oligos for Illumina (Cat# E7335S) kit. Six different indices were mixed for MiSeq analysis, 3 each from the DNA templated or RNA templated assays described above. These included controls (described under “Production of controls for NGS”, two separate identical controls made at different times were used for DNA and RNA templated assays), and products derived from 0.5 or 6 mM MgCl_2_ assays from both the DNA and RNA assays.

### Analysis of NGS results

The basic approach used for analysis of mutations is described in [25] and is broadly termed single strand consensus sequencing (SSCS). Briefly, MiSeq analysis produced ~ 1.6 x 10^7^ individual reads from the 6 different conditions (see above) in the analysis. Reads that differed by more than 5 nucleotides from the predicted length were excluded as the methodologies used to isolate DNA and PCR products (i.e., excision from gels based on size) should have eliminated products that were significantly longer or shorter than “full-length” and these products were therefore assumed to be derived from PCR errors or sequencing errors. Consequently, this approach does not account for long insertion or deletion mutations. However, insertions or deletions of a few nucleotides were scored. After applying various filters to exclude truncated or hypermutated sequences (individual reads with more than 10 errors were excluded), recovered reads were grouped based on the 14 random nucleotide barcode sequence (see “Production of RT DNA products for NGS” above). Reads with the same barcode were assumed to be derived from a single RT extension event. Only barcodes recovered 5 or more times [25] were evaluated and are referred to as a “tag family” in the data. Using 3 rather than 5 as the lower limit for recovered sequences with identical tags did not significantly affect the analysis. On average, there were ~ 230,000 tag families recovered for each condition. The sequence region analyzed was 155 nucleotides in length (see Fig. 1) which yielded an average number of nucleotides evaluated for a specific condition for the RNA templated system (2 rounds of RT synthesis) of ~ 7 x 10^7^ (230,000 x 155 x 2), and ~ 3.5 x 10^7^ (230,000 x 155) for the DNA templated system (1 round of RT synthesis). In order for a particular tag family to register a mutation, the same error had to occur at the same nucleotide position in all of the recovered reads. As expected, based on the error rate of HIV RT, most evaluated tag families had no registered mutations. The MiSeq analysis was conducted twice using independent experiments. Similar results were obtained in the two separate experiments (see Results). To evaluate mutations in the sequence data, we wrote a versatile program that allowed analysis of several parameters including potential errors derived from the MiSeq sequencing process and RT-derived error. This program, along with all the data from these experiments is available from 3 repositories: (1) output logs and some information regarding the process is at: https://github.com/abelew/error_rate_quantification; (2) software which handles the reading of the raw data and creation of the initial tables of mutants per sample is at: https://github.com/abelew/errrt; and (3) postprocessing software at: https://github.com/abelew/Rerrrt/.

## Supporting information

Supplemental Wang et al

## Acknowledgements

We thank Drs. Stefan Sarafianos (Emory University), Eddy Arnold (Rutgers University), and Stephen Hughes (National Institutes of Health) for supplying RTs or RT plasmid clones.

