## Supplemental Wang et al for "Physiological Magnesium Concentrations Increase Fidelity of Diverse Reverse Transcriptases from HIV-1, HIV-2, and Foamy Virus, but not MuLV or AMV"

**by**

**Ruofan Wang[1,3], Ashton T. Belew[1], Vasudevan Achuthan[1,4], Najib El Sayed[1,2], and Jeffrey J. DeStefano[1,2,#]**

Keywords: reverse transcriptase, fidelity, mutation rate, magnesium, retrovirus

[1]Department of Cell Biology and Molecular Genetics, Bioscience Research Building, University of Maryland, College Park, Maryland, USA 20742

[2]Maryland Pathogen Research Institute, College Park, Maryland, USA

[3]Current address: Vigene Biosciences, Rockville Maryland, USA

[4]Current address: CRISPR Therapeutics, Cambridge, Massachusetts, USA

### Table S1: Data from blue/white colony screening (α-complementation assay) of various reverse transcriptases (RT)

| [$Mg^{2+}$] and RT | [a]Colony mutation freqeuncy ((white+faint blue)/total) | [b]Colony mutation frequency - BKG (relative to 0.5 mM HIV-1 wt) | [c]*p* value ($p$<0.05) vs. same enzyme at 0.5 mM $Mg^{2+}$ | [c]*p* value ($p$<0.05) vs. HIV-1 wt at same [$Mg^{2+}$] |
|---|---|---|---|---|
| | | **[d]RNA→DNA→DNA assay** | | |
| Control (BKG) | 16/18,371 (0.00087) | | | |
| 0.25 mM HIV-1 wt | 49/13,180 (0.00372) | 0.00285 (0.82) | 0.505255* | NA |
| 0.5 mM HIV-1 wt | 29/6,672 (0.00435) | 0.00348 (1) | NA | NA |
| 1 mM HIV-1 wt | 36/6,608 (0.00545) | 0.00458 (1.3) | 0.365522* | NA |
| 2 mM HIV-1 wt | 65/9,816 (0.00662) | 0.00575 (1.7) | 0.058194* | NA |
| 6 mM HIV-1 wt | 79/9,305 (0.00849) | 0.00762 (2.2) | 0.001736 | NA |
| 12 mM HIV-1 wt | 87/11,348 (0.00767) | 0.00680 (2.0) | 0.007477 | NA |
| 0.5 mM HIV-1 A/E wt | 15/3,688 (0.00407) | 0.00320 (0.92) | NA | 0.834904* |
| 6 mM HIV-1 A/E wt | 40/4,965 (0.00806) | (0.00719) (2.1) | 0.021726 | 0.787848* |
| 12 mM HIV-1 A/E wt | 32/4,005 (0.00799) | (0.00712) (2.0) | 0.028338 | 0.842173* |
| 0.5 mM PFV wt | 27/11,052 (0.00244) | (0.00157) (0.45) | NA | 0.029231 |
| 6 mM PFV wt | 30/5,997 (0.00500) | (0.00413) (1.2) | 0.005881 | 0.012866 |
| 12 mM PFV wt | 36/6,431 (0.00560) | (0.00473) (1.4) | 0.000827 | 0.112193* |
| 0.5 mM HIV-2 wt | 39/12,232 (0.00319) | 0.00232 (0.67) | NA | 0.205403* |
| 6 mM HIV-2 wt | 50/7,576 (0.00660) | 0.00573 (1.6) | 0.000517 | 0.163867* |
| 12 mM HIV-2 wt | 69/7,858 (0.00880) | 0.00793 (2.3) | <0.00001 | 0.453543* |
| 0.5 mM K103N HIV-1 | 22/5,819 (0.00378) | 0.00291 (0.84) | NA | 0.617957* |
| 6 mM K103N HIV-1 | 39/5,321 (0.00733) | 0.00646 (1.9) | 0.011695 | 0.453918* |
| 0.5 mM AZT[R] HIV-1 | 11/3,000 (0.00367) | 0.00280 (0.80) | NA | 0.631232* |
| 6 mM AZT[R] HIV-1 | 28/2,953 (0.00948) | 0.00861 (2.5) | 0.005736 | 0.616860* |
| 0.5 mM K65R HIV-1 | 26/8,960 (0.00290) | 0.00203 (0.58) | NA | 0.132730* |
| 6 mM K65R HIV-1 | 40/7,663 (0.00522) | 0.00435 (1.3) | 0.018301 | 0.011638 |
| 0.5 mM M184V HIV-1 | 40/11,141 (0.00359) | 0.00297 (0.85) | NA | 0.433469* |
| 6 mM M184V HIV-1 | 101/8,305 (0.01216) | 0.01129 (3.2) | <0.00001 | 0.016707 |
| 0.5 mM Q151M HIV-1 | 9/2,588 (0.00349) | 0.00262 (0.75) | NA | 0.558781* |
| 6 mM Q151 M HIV-1 | 19/2,462 (0.00772) | 0.00685 (2.0) | 0.043717 | 0.709775* |
| 0.5 mM MuLV | 19/4,940 (0.00385) | 0.00298 (0.86) | NA | 0.679027* |
| 6 mM MuLV | 12/2,836 (0.00423) | 0.00336 (0.97) | 0.7961* | 0.022173 |
| 0.5 mM AMV | 31/6,120 (0.00507) | 0.00420 (1.2) | NA | 0.554112* |
| 6 mM AMV | 34/5,497 (0.00619) | 0.00532 (1.5) | 0.421766* | 0.122281* |
| | | **[d]DNA→DNA assay** | | |
| Control HIV-1 (BKG) | 10/11,078 (0.00090) | | | |
| 0.5 mM HIV-1 wt DNA | 8/4,343 (0.00189) | 0.00099 (1) | NA | NA |
| 6 mM HIV-1 wt DNA | 36/7,261 (0.00496) | 0.00406 (4.1) | 0.00844 | NA |
| Control MuLV (BKG) | 3/5,198 (0.00058) | | | |
| 0.5 mM MuLV DNA | 14/7,256 (0.00193) | 0.00135 (1.4) | NA | 0.916775* |
| 6 mM MuLV DNA | 8/5,928 (0.00135) | 0.00077 (0.8) | 0.417814* | 0.001538 |

a- Approximately 200 colonies per 100 mm plate were visually scored. Total = white + faint blue + blue. Questionable faint blue colonies were replated with blue colonies to further access the color.
b- Colony mutation frequency was the (white+faint blue)/total values in column 2 and the BKG is from the "Control (BKG)" in column 2. For the RNA→DNA→DNA assay, BKG controls were made from PCR of plasmid sequences so they do not represent the "real" assay BKG. For DNA→DNA assays, BKG controls were from PCR of the starting DNA template and therefore represent the "true" BKG for the assay. See Methods and Results for an explanation of the BKG.
c-*p* values were calculated using Chi Square analysis and a significance level of $p < 0.05$. Values less than 0.05 were considered significant.
d- The RNA→DNA→DNA and DNA→DNA assays are described under Methods and Fig. 1 (main manuscript). The former includes two RT synthesis steps and the latter just 1. The former also starts with an RNA template made from T3 RNA polymerase. Therefore, more mutations would be expected in the RNA→DNA→DNA assay with the same reaction conditions.
*Value considered not statistically significant (>0.05) based on Chi-squared statistical analysis.

### Table S2. Data from NGS experiment 2

| Condition[a] | Control DNA and RNA | DNA 0.5 mM $Mg^{2+}$ | DNA 6 mM $Mg^{2+}$ | RNA 0.5 mM $Mg^{2+}$ | RNA 6 mM $Mg^{2+}$ |
|---|---|---|---|---|---|
| **Total Indices[b]** | 303,378 | 301,642 | 185,208 | 149,329 | 388,824 |
| **Total nucleotides[c]** | $4.702 \times 10^7$ | $4.675 \times 10^7$ | $2.871 \times 10^7$ | $4.629 \times 10^7$ | $1.205 \times 10^8$ |
| **Substitution Type** | **Number of recovered substitutions (substitution frequency per nucleotide x $10^6$)[d]** | | | | |
| A>C | 729 (15.50) | 1091 (23.33) | 2001 (69.70) | 964 (20.82) | 3939 (32.68) |
| A>G | 367 (7.805) | 2073 (44.34) | 1862 (64.86) | 909 (19.64) | 3761 (31.20) |
| A>T | 456 (9.697) | 721 (9.753) | 1811 (15.88) | 790 (9.851) | 3954 (32.81) |
| *C>A | 6669 (141.8) | 6595 (141.1) | 10448 (365.3) | 4704 (101.6) | 18361 (152.3) |
| C>G | 613 (13.04) | 2083 (44.55) | 4178 (145.5) | 1282 (27.69) | 12996 (107.8) |
| C>T | 1605 (34.13) | 3252 (69.55) | 14521 (505.8) | 3029 (65.43) | 26741 (221.9) |
| G>A | 602 (12.80) | 8091 (173.1) | 13951 (486.0) | 2547 (55.02) | 20808 (172.6) |
| G>C | 271 (5.763) | 368 (7.871) | 523 (18.22) | 185 (3.996) | 1034 (8.578) |
| *G>T | 16002 (340.3) | 11235 (240.3) | 8044 (280.2) | 5894 (127.3) | 30484 (252.9) |
| T>A | 745 (15.84) | 1121 (23.98) | 2170 (75.59) | 1059 (22.88) | 3569 (29.61) |
| T>C | 251 (5.338) | 600 (12.83) | 7609 (265.1) | 991 (21.41) | 7300 (60.56) |
| T>G | 897 (19.08) | 1269 (27.14) | 2041 (71.10) | 843 (18.21) | 3545 (29.41) |
| **Insertion Type** | **Number of recovered insertions (insertion frequency per nucleotide x $10^6$)[d]** | | | | |
| A | 0 | 0 | 151 (5.359) | 266 (5.746) | 204 (1.693) |
| C | 0 | 0 | 148 (5.155) | 58 (1.253) | 138 (1.145) |
| G | 0 | 0 | 16 (0.5573) | 0 | 24 (0.1992) |
| T | 0 | 19 (0.4064) | 1511 (52.63) | 1357 (29.32) | 1119 (9.286) |
| **Deletion Type** | **Number of recovered deletions (deletion frequency per nucleotide x $10^6$)[d]** | | | | |
| A | 52 (1.106) | 124 (2.652) | 374 (13.03) | 144 (3.112) | 669 (5.552) |
| C | 45 (0.9570) | 183 (3.930) | 914 (31.84) | 501 (10.82) | 2087 (17.32) |
| G | 45 (0.9570) | 48 (1.031) | 199 (6.931) | 131 (2.830) | 640 (5.311) |
| T | 61 (1.297) | 113 (2.426) | 787 (27.41) | 246 (5.314) | 1319 (10.95) |
| **Mutation Type** | **Percent of various types of mutations for each condition (mutation frequency x $10^4$)[e]** | | | | |
| *Substitutions | | 98.0 (2.975) | 91.9 (15.78) | 70.4 (1.286) | 92.6 (5.883) |
| Insertions | | 0.1 (0.0030) | 3.7 (0.6353) | 19.9 (0.3637) | 1.9 (0.1207) |
| Deletions | | 1.9 (0.0577) | 4.4 (0.7555) | 9.7 (0.1772) | 5.5 (0.3494) |
| Overall mutation frequency | | $3.036 \times 10^{-4}$ | $1.717 \times 10^{-3}$ | $1.827 \times 10^{-4}$ | $6.353 \times 10^{-4}$ |

a- Refer to Material and Methods for details. DNA assays included 1 round of RT DNA-template-directed DNA synthesis while RNA assays include an RT RNA-template-directed round and a second RT DNA-template-directed DNA synthesis round.
b- Indices are defined in Materials and Methods. To qualify as an “index”, the same 14-nucleotide barcode had to be recovered in 5 or more reads in NGS. Specific mutations were recorded when they appeared at a specific position in all the reads that constituted a particular index.
c- Total number of RT-directed nucleotides catalysis events evaluated in the assay. For DNA templated assay, this was the number of indices x 155 nucleotide positions (see Fig. 1, main manuscript). For RNA templated assays, the indices number was multiplied by 310 because this assay had an additional round of RT-directed synthesis (see Materials and Methods).
d- Insertion frequency per nucleotide was calculated by dividing the raw number of mutations of the specified type by the total number of nucleotides that were assessed for that condition. Unlike the raw number of mutations, this value allows a direct relative comparison of the data in the Table with higher numbers representing a greater frequency for that mutation. Note that the DNA and RNA templated assays cannot be directly compare with this number the as the templates was synthesized by Q5 DNA polymerase or T3 RNA polymerase in the DNA and RNA assays, respectively. The Q5 synthesis product was used as the Control for both assays. Given the very low mutation rate of Q5, it is likely the RNA assays contain a significant proportion of mutations derived from T3 RNA polymerase that are not accounted for by the Control (see Results for further discussion).
e- Values were determined after subtracting background Control values from each mutation type using the frequency per nucleotide data above. These values were then summed for each of the 3 categories (i.e. substitutions (S), insertions (I), and deletions (D)). The percent values listed were calculated by dividing individual values by the sum of the 3 categories and multiplying by 100 (e.g. for Substitutions: (S/(S+I+D)) x 100). Mutation frequencies in parentheses were the individual S, I, and D values and the Overall Mutation Frequency was S+I+D for each category.
*Due to high numbers in the Controls, C>A and G>T substitutions were excluded from data calculations (see Results).

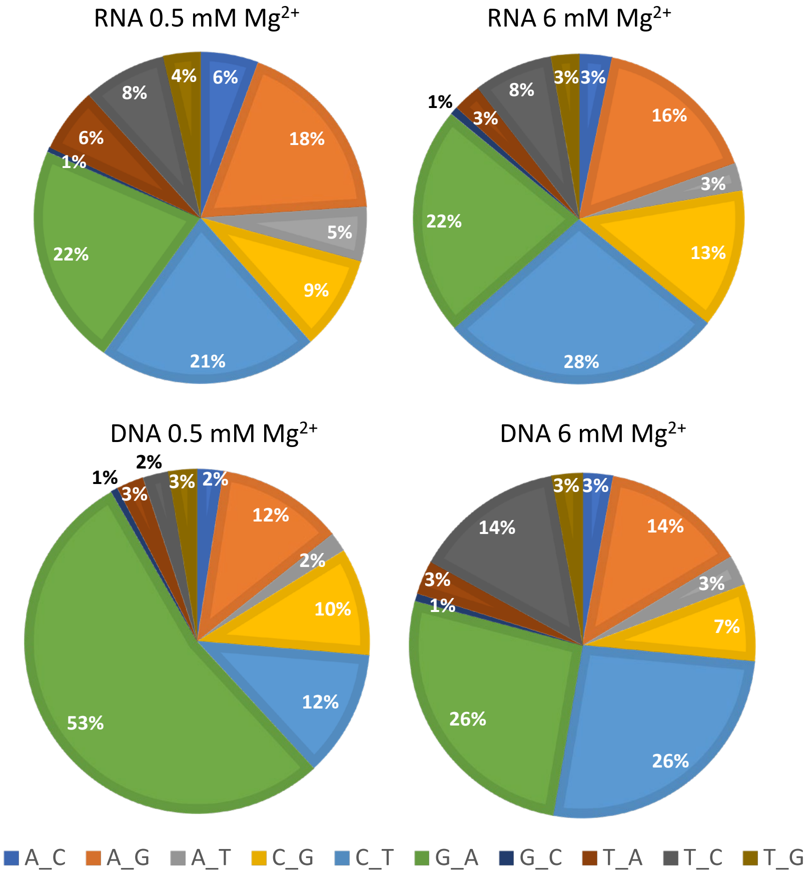

Figure S1. Proportion of various substitution mutations in the NGS assay for Exp. 2. The proportions of substitution mutations recovered using the indicated conditions in the NGS assay are shown. G>T and C>A mutations are excluded from the analysis (see Results). See Table S2 for more information on the mutation rates for the various mutation types.

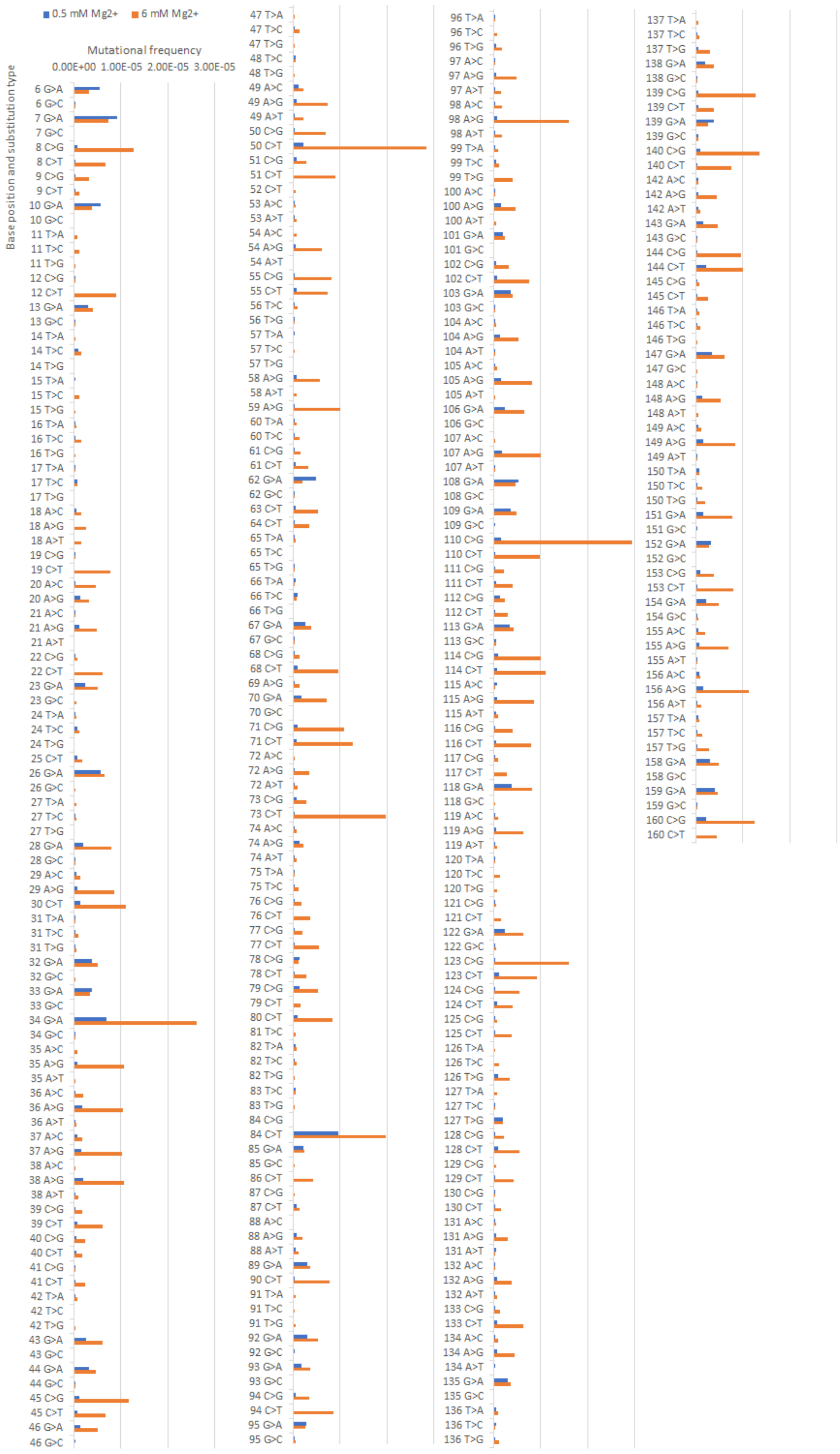

Figure S2. Substitution mutation frequencies using the DNA templated system in the NGS assay. The mutation frequency per nucleotide at specific positions (see Table 1 in main text) for specific types of recovered substitution mutations at the two different $Mg^{2+}$ concentrations (blue, 0.5 mM and orange 6 mM $Mg^{2+}$) is plotted. Mutation frequencies are for the specific mutation type and position and were calculated after subtraction of background rates at the same template position. See Table 1 for the total number of nucleotides scored for each condition.
